# A quantitative approach to assess similarity between dives under the Bühlmann decompression algorithm constraints

**DOI:** 10.64898/2026.09.16.752047

**Authors:** Joshua B. Currens, Frauke Tillmans, Virginie Papadopoulou

## Abstract

Undersea and high-altitude operations regularly put operators at risk for decompression sickness (DCS), which is attributed to bubble formation and growth during decompression. DCS mitigation strategies are validated with empirical testing, often conducted in hyperbaric chambers, which provide precise control of the pressure-time profile. A degree of variability for study outcome is anticipated in both decompression sickness (DCS) and venous gas emboli (VGE) endpoints. Experimentation in a chamber offers precision, but is not always practical or operationally relevant, leading researchers to use open water for the dive exposure. Dive profile consistency may vary by diver skill level and environmental conditions, potentially introducing dive profile deviations, which may be a factor of individual variability. There does not exist a quantitative framework for evaluating dive profile consistency and quantifying deviations in open water studies. In this work, we evaluate existing dive profile metrics and propose a workflow to estimate dive profile deviations using the Bühlmann dissolved gas phase model. The dive profile metrics included depth and time-based calculations, as well as surface gradient factor, degree-of-conservatism and the root mean square error of the Bühlmann tissue compartments. We developed a set of simulated dive profiles and obtained a set of 40 open water dives previously collected by certified divers, both of which were used to evaluate performance of the dive profile metrics. Depth and time-based metrics offered sensitivity to varying depths and time durations of the dive profiles, respectively. For the field dives, we found the Bühlmann root mean square error metric performed best at identifying the deviation profile, followed by pressure root time. These evaluations demonstrate the utility and limitations of previously published dive profile metrics and emphasize the importance of accounting for dive profile consistency in open water decompression trials.

**New & Noteworthy:** Inter-individual variability in response to decompression stress has been documented for participants completing the same dive exposure, in either a hyperbaric chamber or in open water. Dives in open water are not afforded the same pressure-time control as those in a hyperbaric chamber. We developed a pipeline using an established dissolved gas phase model to quantify depth-time profile deviation for open water human diver studies. We evaluated our workflow against pre-existing metrics and found inconsistencies. Dive profile variability should be considered in open-water decompression studies.

## Introduction

Undersea operation and exploration can expose the individual to an elevated risk of decompression sickness (DCS) during and after return to atmospheric pressure. DCS occurs after a reduction of pressure causes excess bubble formation in the body, inducing a range of effects from skin rash to paralysis or even death (1, 2). Current DCS mitigation strategies are imperfect, and most documented cases recreationally today are found to occur despite following safe diving guidelines (3). For decompression table development and intervention evaluation, researchers use empirical testing to confirm the safety of a profile or efficacy of a pre-dive intervention (4, 5), either measuring DCS rate or relying on venous gas emboli (VGE) presence as a surrogate outcome (6, 7). Many previous experiments have been conducted in hyperbaric chambers, where the temperature and pressure can be precisely controlled (8–18). Other studies have been conducted in a field environment (such as a pool, lake, or ocean), with less control of the exposure for each diver (19–21). Variability of DCS occurrence and VGE presence has been reported in the hyperbaric chamber setting (8, 18, 22) and open water dives (3, 20, 21); however, the reduced exposure control in the open water context may confound the analysis. Furthermore, DCS has been reported following dive schedules that individuals had previously completed without adverse effects (3, 22). However, similarity between exposures is difficult to assess, and dive profile deviations may have contributed to DCS occurrence. Previous researchers have sought to address the challenge of dive profile consistency by providing an in-water guide for each dive team to ensure compliance (23), yet this still offers less control than the chamber setting. Focusing on studies conducted in open water or a pool, several indicate a pre-determined dive plan with decompression parameters to guide the return to surface (19–21), which relies on participant skill and adherence to computer guidance. Monitoring depth and time with a dive computer also allows for visual comparison against the prescribed profile for deviations (19, 24–30), providing some information but this does not provide a quantitative threshold or standard. For example, Dujic *et al*. evaluated the average depth of dives to estimate similarity between a control and intervention group (31). There does not exist a standard method of dive profile monitoring for open water exposures.

Multiple approaches to decompression modeling have been previously developed to minimize the DCS risk of a given dive profile and can be broadly categorized as either deterministic or probabilistic models; however, no method has been established to quantitatively assess how minor deviations from a planned dive profile affect DCS risk. This gap is particularly relevant for studies investigating inter- and intra-subject variability in DCS or venous gas emboli (VGE), which often treat nominally identical open-water dives as equivalent exposures rather than evaluating observed outcomes against the variability in decompression risk predicted from actual dive profile differences.

Deterministic algorithms rely solely on depth and time calculations to set a depth ceiling that, if exceeded, would result in excessive bubble development and an unacceptable risk of DCS (32). The Bühlmann 16-compartment algorithm with gradient factors is one of the most prevalent computer implementations for divers intentionally conducting dives with a decompression obligation (33, 34). Analysis of inert gas theoretical tissue tension derived from the Bühlmann compartments has been discussed in the rat model for predicting DCS status (35). In 1998, Baker described the utility of gradient factors to further customize the level of conservatism to the Buhlmann model (33), subsequently allowing the calculation of a surface gradient factor, representing the maximum level of critical supersaturation at the end of dive. The surface gradient factor has been evaluated for relationship to DCS at scale in a population database (36). However, the Bühlmann framework has not been evaluated for identifying individual deviations while attempting a prescribed open-water dive profile. In 1952, Hempleman *et al*. proposed a decompression limit based on maximum pressure and square root of time (Prt) to approximate the total inert gas charge for a given dive, which could be used to approximate depth limits (37). In the past, Shields *et al*. demonstrated that the Hempleman Prt approach could be used to calculate a generalized metric that correlated with DCS rates in commercial diving operations (38), but this approach does not account for the decompression phase of the dive. Separately, a probabilistic method to DCS mitigation has been described and implemented using a log likelihood model that incorporates depth, time, and has been trained using previously conducted experiments (39). However, both Prt by Shields *et al*. (38) and the probabilistic approach described by Weathersby *et al*. (39) are intended for evaluation of large profile differences and may not be sensitive to the individual differences of divers attempting to follow the same profile.

To date, a framework does not exist for quantitatively evaluating the deviation of experimental attempts at a prescribed dive profile, which may relate to VGE or DCS risk variability in open water experimental dives. Deterministic decompression algorithms, such as Bühlmann (ZHL-16C), offer insight to theoretical inert gas kinetics during a dive, which may be leveraged to identify slight deviations. Development of a method for comparing open water dive profiles would allow for evaluation of the total deviation for a given dive, which may be used to validate the consistency of a set of dives. Additionally, a method to estimate the deviation of a given dive against the prescribed exposure could help researchers determine if the variability in VGE, or subsequent DCS risk, is related to small differences in the dive exposure. In this paper, we will apply the Bühlmann workflow to evaluate sensitivity to prescribed profile deviations and compare against other depth and time-based metrics, such as Prt and gradient factors.

## Methods

### Ethics Statement

The open-water dive profiles used for illustration in this study were obtained from an ongoing study conducted by the Divers Alert Network and the University of North Carolina at Chapel Hill. The study received Institutional Review Board (IRB) approval (DAN IRB No.:029- 21-27; UNC IRB No: 21-3299), and all participants provided informed consent prior to enrollment. The present analysis used only previously acquired, de-identified depth–time profile data; no additional participant data were accessed or analyzed.

### Depth and Time Summary Metrics

Dive status is typically monitored using a wrist mounted computer that tracks depth and time at regular intervals. Summary metrics were identified and calculated including maximum depth, average depth, bottom time, and total time. The bottom time was defined as the time from start of dive until the first instance the diver exceeded 2 meters shallower than the maximum depth of the dive. The average and maximum instantaneous ascent rate (*v_inst_*) calculated throughout the decompression phase of the dive can be used to describe all moments of pressure decrease using Eqn. 1. Area-under-the-curve (*AUC*) of the dive profile was evaluated using a trapezoidal sum across the dive duration (Eqn. 2). For Eqns. 1&2, depth (*d*) was measured in meters and time (*t*) in minutes.

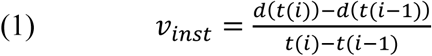

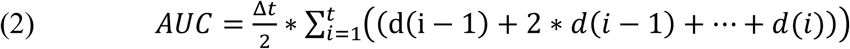

### Bühlmann Algorithm Implementation

The Bühlmann ZHL-16C decompression model was replicated in MATLAB (The MathWorks, Inc., Natick, MA, USA), allowing for estimation of theoretical nitrogen uptake for each of the 16 compartments (40). The ZHL-16C algorithm is based on the Haldane approach of limiting tissue supersaturation and was selected for its wide integration in commercial dive computers (34). ZHL-16C is described in detail elsewhere (40), for brevity only key details are reproduced. Each of the 16 compartments represents a different theoretical tissue type, with varying half-times of inert gas absorption and release. The 16 compartment halftime values for nitrogen can be found in Table 1, conversion to helium half-times can be obtained by dividing the nitrogen values by 2.65 to account for differences in gas kinetics (40). Figure 1 presents an example dive profile with the theoretical nitrogen partial pressure for the 16 different tissue compartments.

**Figure 1.**
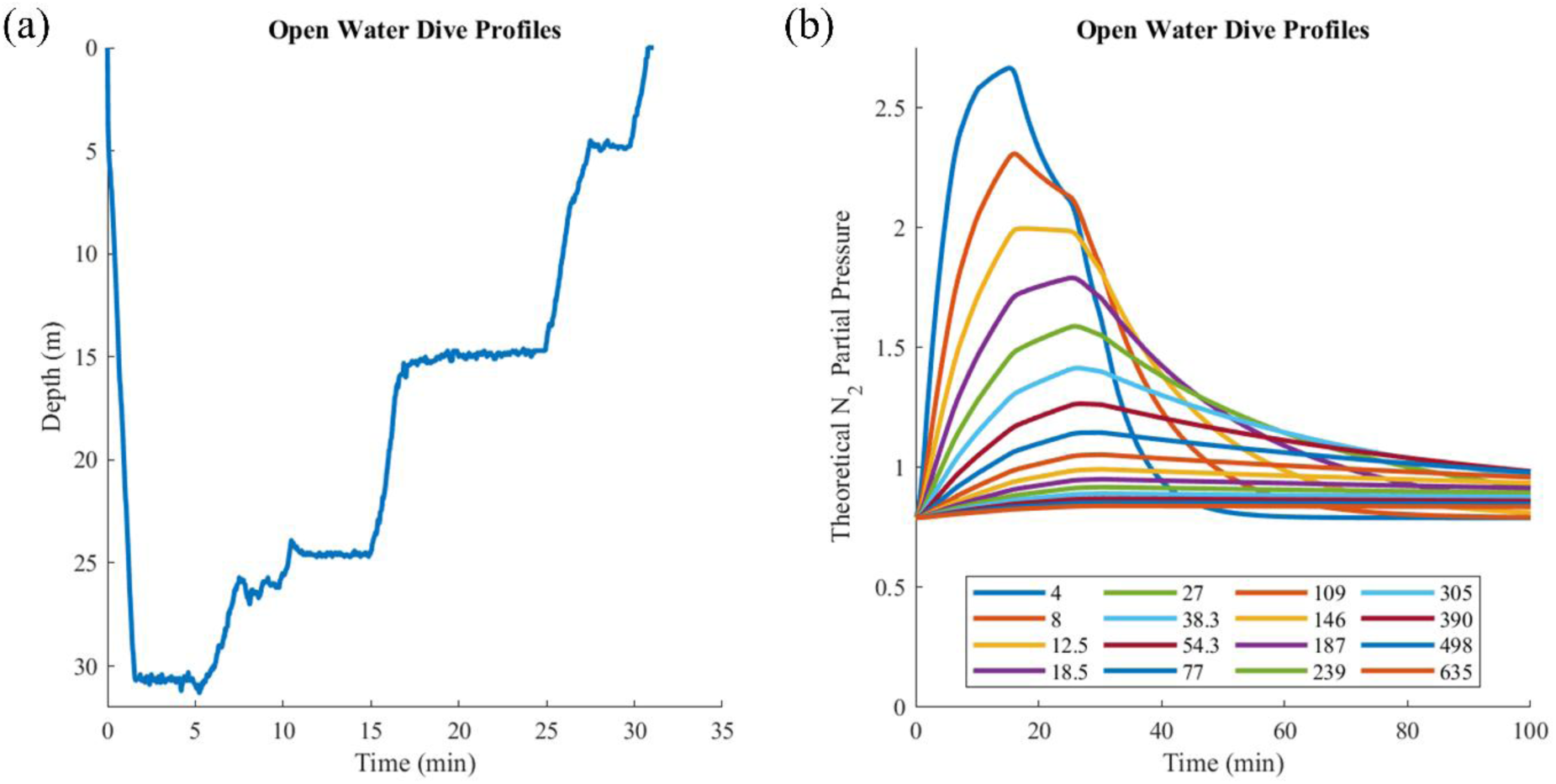
Example dive profile and Bühlmann implementation. (a) Dive profile obtained from dive computer and the (b) 16 theoretical tissue compartments during the dive and for 60 minutes after surfacing.

**Table 1.** Bühlmann Nitrogen Half-times.

| Compartment | 1 | 2 | 3 | 4 | 5 | 6 | 7 | 8 | 9 | 10 | 11 | 12 | 13 | 14 | 15 | 16 |
| --- | --- | --- | --- | --- | --- | --- | --- | --- | --- | --- | --- | --- | --- | --- | --- | --- |
| N<sub>2</sub> Halftimes<br>(h<sub>N2</sub>) | 4 | 8 | 12.5 | 18.5 | 27 | 38.3 | 54.3 | 77 | 109 | 146 | 187 | 239 | 305 | 390 | 498 | 635 |

Decompression (or returning to surface) under the deterministic ZHL-16C model is conducted by estimating the inert gas tension in each of the 16 compartments (Eqn. 3).

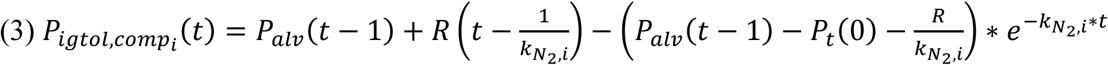

Where, the total inert gas partial pressure (*P_igtol_*) per compartment is calculated at a given ambient pressure (*P_t_*) based on the alveolar partial pressure (*P_alv_*) and rate of change (*R*), gas specific rate constants per compartment (*k_N2,i_*) for a given time (*t*) (Eqn. 3). The rate constant (*k*) is based on the half-time values (represented by *t_N2,i_*) for the inert gas in the breathing mix, providing a value for each tissue compartment (Eqn. 4).

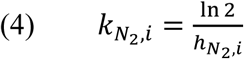

Next, the ambient pressure ceiling (*P_amb,ceiling_*) per compartment is defined as the lowest tolerable ambient pressure the individual could sustain without exceeding critical tissue supersaturation levels (Eqn. 5).

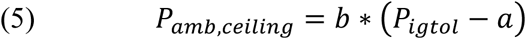

Calculation of the inert gas tissue tension limit is unique for each compartment and relies on variables *a* and *b* found through Eqns. 6 and 7, respectively.

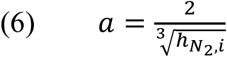

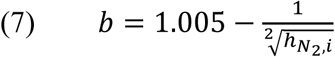

Where, *h_N2,i_* denotes the compartment half time for nitrogen. The compartment closest to the relative critical supersaturation is identified as the ‘leading compartment’, which dictates the shallowest depth a diver can reach without exceeding the critical inert gas tissue tension threshold. A pressure change that exceeds the ceiling of the current time is considered to breach the safe level of tissue supersaturation and increase the likelihood of *de novo* bubble formation as well as risk of DCS. Equations can be modified for helium breathing gas mixtures by replacing the nitrogen half-times with the helium values.

### Bühlmann Algorithm-Based Analysis

To estimate dive profile risk of exceeding the Bühlmann model constraints, a degree of conservatism (DoC) metric was defined (41). The DoC was taken as the ratio of the instantaneous pressure to the pressure ceiling calculated at the previous instance, which is applied for all 16 compartments (Eqn. 8). This approach provides a DoC value for all 16 compartments at every recorded time point, using the dive computer sampling interval. The minimum DoC >1 value across the dive represents the closest an individual came to exceeding the pressure ceiling, and any DoC value less than 1 indicates that a ceiling was surpassed. The overall minimum DoC and average minimum DoC were calculated throughout each dive.

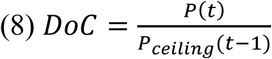

Where, *P(t)* is the ambient pressure at time *t* and *Pceiling(t-1)* is the previous pressure ceiling obtained in Eqn. 5. Separately, the inert gas tissue tension values per compartment were compared. When a diver has attempted to follow a prescribed profile, the root-mean-square-error (RMSE) between each can be evaluated to directly compare each of the 16 compartments. The average RMSE (*pp_N2,RMSE_*) over the 16 compartments was obtained to describe the deviation between the test profile and the reference dive (Eqn. 9).

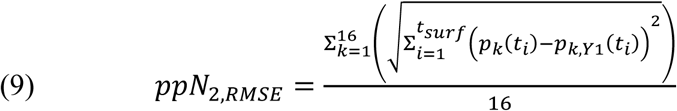

Where, *t_surf_* is the time of surfacing, *P_k_(ti)* is the inert gas partial pressure per compartment k at time ti for the test profile, and P*_k,Y1(_t_i_)* is the inert gas partial pressure per compartment k at time ti for the reference profile.

### Previously Published Dive Metrics

To evaluate the prevalence of DCS in a dataset of commercial dive profiles, Sheilds *et al*. utilized an index based on maximum pressure and bottom time (Prt) (38), derived from Hempleman *et al*.’s theoretical limit (37). While simple, this approach provides an approximate threshold to prevent an unacceptably high risk of DCS, based on the approximate decompression obligation required for the Prt (37). We implemented this calculation by identifying the maximum pressure reached throughout the dive. For computational calculations, the bottom time was defined as the time when the individual reached 2 meters shallower than maximum depth during the return to surface, and verified for each dive. The maximum pressure (*P_max_*) times the square root of bottom time (*T_b_*) provides Prt, calculated using Eqn. 10 (37).

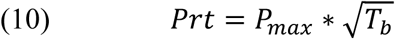

Gradient factors (GF) were first described for use in the Bühlmann algorithm by Baker in 1998 to factor in additional conservatism for technical dives with decompression obligation (33). The GF development allowed for granular threshold adjustment of the pressure ceiling to be more conservative than the critical M-values. GF values per compartment were calculated throughout each dive using Eqn. 11, representing the current percentage of critical supersaturation per compartment. The maximum GF at time of surfacing (GF_surf_) across the 16 compartments represents the percent of maximum tolerable tissue supersaturation that occurred when the diver completed the dive.

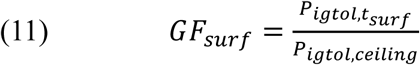

Where, *P_igtol,tsurf_* is the inert gas partial pressure at the time of surfacing and *P_igtol,ceiling_* is the inert gas partial pressure ceiling for an ambient pressure of 1 atmosphere.

### Simulated Dive Profiles

To evaluate the dive metrics described thus far, several test profiles were simulated in MATLAB. One set of simulated profiles range in depth (depth-variable) within the limits of air diving, spanning 10 to 70 meters in depth at 10-meter intervals (Fig. 2a). The 50-, 60- and 70-meter dives would require staged decompression in practice, and the latter profile is considered an ‘exceptional exposure’ by the US Navy Diving Manual (42). A second set (time-variable) maintained a 30- meter maximum depth with varying bottom times from 5 to 75 minutes at 10-minute intervals (Fig. 2b). For the time-variable dives, bottom times of 30 minutes and greater require decompression by the US Navy Diving Manual (42), and the 75-minute profile is considered an ‘exceptional exposure’. Based on recreational diving guidelines, descent and ascent rates for all simulated dives were set to 18 and 8 meters per minute, respectively. Across both sets of simulated profiles, the shallow and shorter dives were within recreational diving limits.

**Figure 2.**
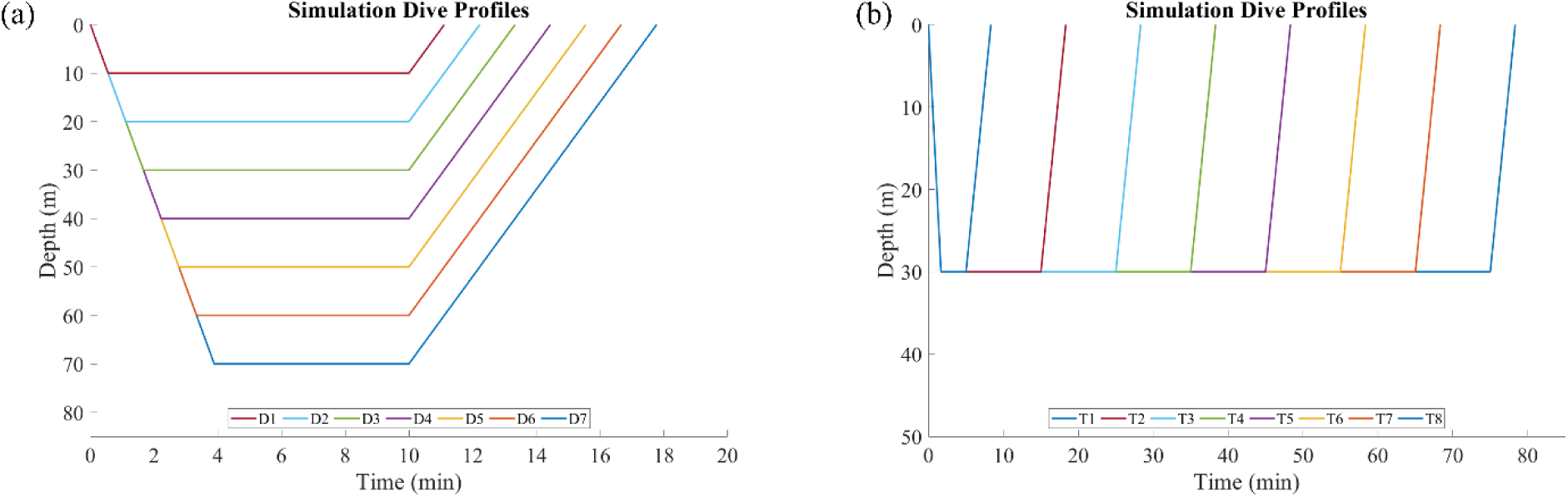
Simulation dive profiles with varying maximum depths (a) and bottom times (b).

For each profile, the following depth and time-based metrics were calculated: average depth, maximum depth, total dive time, bottom time, average ascent rate, maximum ascent rate, and area-under-curve for depth-time. Additionally, Prt and Bühlmann-based metrics: average minimum DoC, minimum instantaneous DoC, and GF_surf_ were obtained for the simulated dives. To compare the metrics on a similar scale, the percent difference was taken for the depth- and time-variable profiles separately, with the reference being taken as the average of the profile set.

### Field dive dataset

A total of 40 de-identified real open-water dive profiles were also analyzed. The dive profile was established prior to data collection and included approximately five minutes at the 30-meter maximum depth with a multi-level return to surface for a total run time of 30 minutes. A guideline with directional arrows was maintained to provide visual guidance for the divers. Individual field dives were recorded using a wrist-mounted Teric dive computer (Shearwater Research Inc., Richmond, British Columbia, Canada) with a 0.5 Hz sample rate. The profile data was exported after each dive. Data handling and visualization was conducted in MATLAB. A 10-second moving average of each dive profile was taken to reduce noise due to minor position adjustments.

A subset of three dives was evaluated for differences in DoC overtime. One profile was the average of 39 dive attempts at the prescribed profile to use as a standard (Average Profile) to compare against. Notably, one of the divers reported difficulty with middle ear equilibration and was delayed in reaching the maximum depth, labeled as ‘deviation profile’. The third profile simulated a dive with a 25-meter maximum depth that follows the average profile for the return to surface and is referred to as ‘depth-augmented’ going forward. For this subset, the DoC was calculated for each compartment overtime.

## Results

The depth and time summary metrics, Bühlmann algorithm-based analysis, Prt, and GF_surf_ were calculated for the simulated and field collected dive profiles. Results for the two groups of profiles are presented separately. Values are presented as mean ± standard deviation, unless otherwise noted.

### Simulated Dive Profiles

The depth- and time-variable simulation profiles covered a range of relevant air dive exposures. Differences in average and maximum depth varied the most for the depth-variable profiles (Fig. 3a-b). While the time-variable profiles demonstrated more deviation in total dive time and bottom time (Fig. 3c-d). No differences were identified in average or maximum ascent rate (Fig. 3e-f). The dive AUC was found to increase orderly with both depth and time (Fig. 3g). Time-variable profiles to 30-meter maximum depth, with 25-minute or longer bottom time, had a larger AUC than the 70-meter dive with a 10-minute bottom time (Fig. 3g).

**Figure 3.**
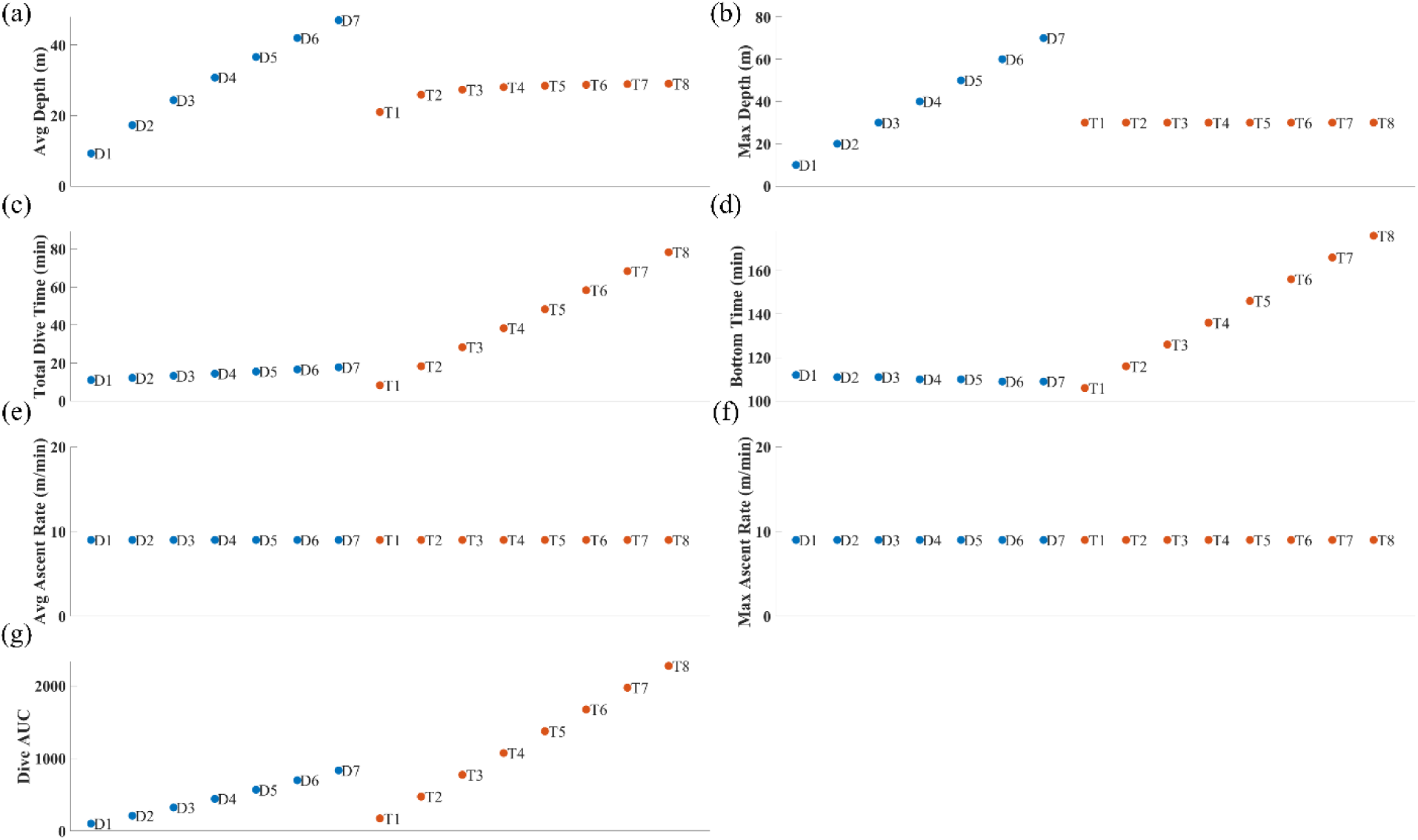
Depth and time-based metrics for simulated dive profiles. (a) Average Depth, (b) Maximum Depth, (c) Total Dive Time, (d) Bottom Time, (e) Average Ascent Rate, (f) Maximum Ascent Rate, and (g) Dive Profile Area-Under-Curve. For all plots, dives are evenly spaced horizontally to prevent overlap.

Analysis of the simulation profiles via the DoC from the Bühlmann framework, GF_surf_, and Prt is presented in Figure 4. The average DoC calculated throughout the return to surface varied orderly with time but not depth (Fig. 4a). Minimum DoC changed inversely with increasing depth and time profiles (Fig. 4b). The maximum GF_surf_ across the 16-compartments increased with both depth and time, with several exceeding a value of 1 (Fig. 4c). Calculations of Prt also demonstrated a positive relationship with depth and time increases (Fig. 4d). Three depth-variable profiles (50-, 60-, and 70-meters) and six time-variable profiles (25 to 75-minutes) had a minimum DoC less than 1 at some time during the return to surface and a GF_surf_ greater than 1, indicating the pressure ceiling was exceeded.

**Figure 4.**
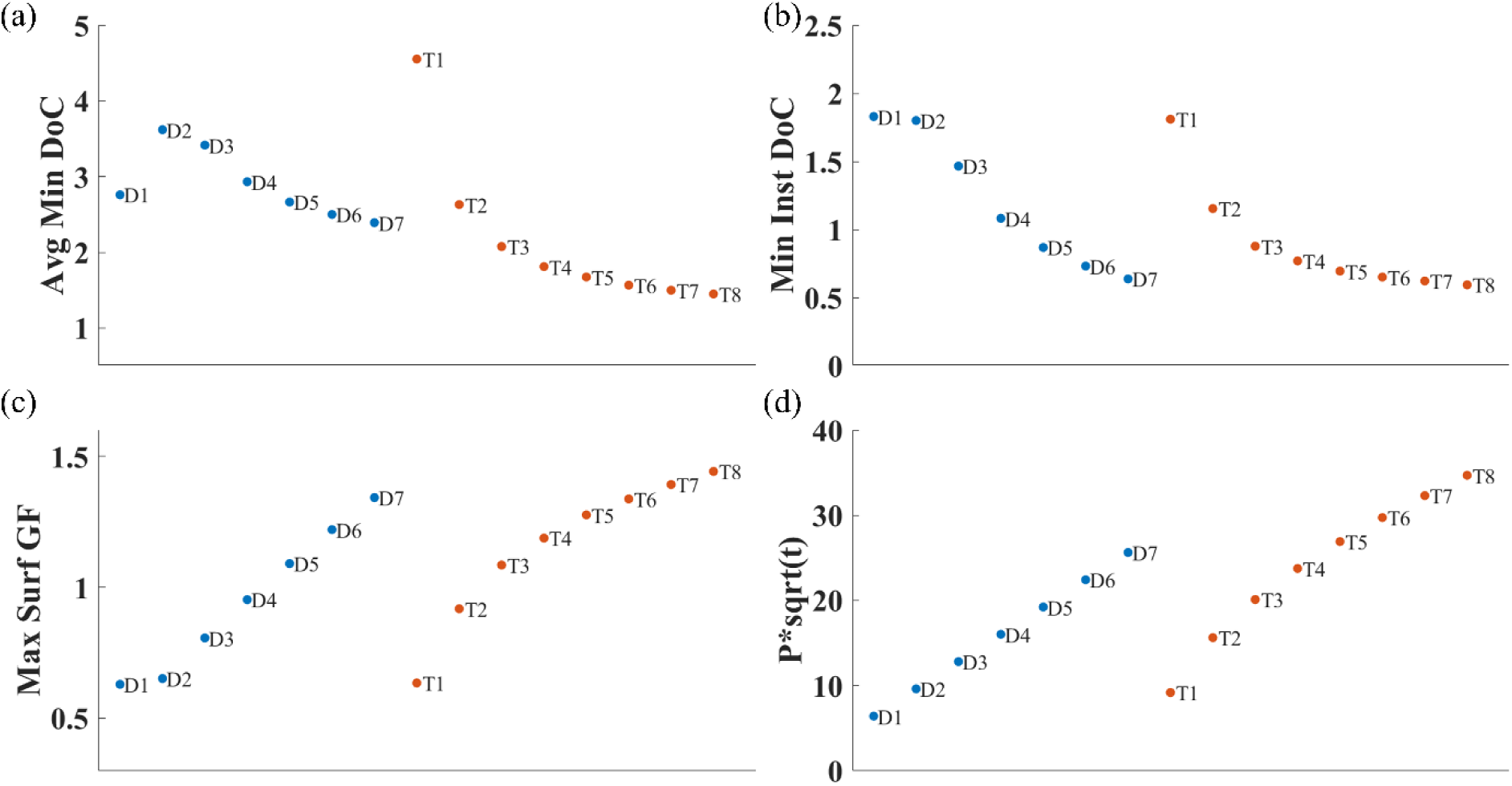
Analysis of simulation dive profiles under Bühlmann workflow and previously established metrics. (a) Average Minimum Degree of Conservatism (DoC), (b) Instantaneous Minimum DoC, (c) Maximum Surface Gradient Factor (GF), and (d) Maximum Pressure root Time (Prt). For all plots, blue and orange represent depth- and time-variable simulation profiles, respectively.

Percent difference for each metric within depth- and time-variable respective datasets is presented in Figure 5. Dive AUC and Prt were found to have a large spread in percent difference for both depth- and time-variable profiles. Average and maximum depth had a wide range of percent difference for the depth-variable profiles, yet minimal change for the time-variable dataset. Comparatively, total dive time and bottom time better differentiated the time-variable dataset, but not the depth-variable profiles.

**Figure 5.**
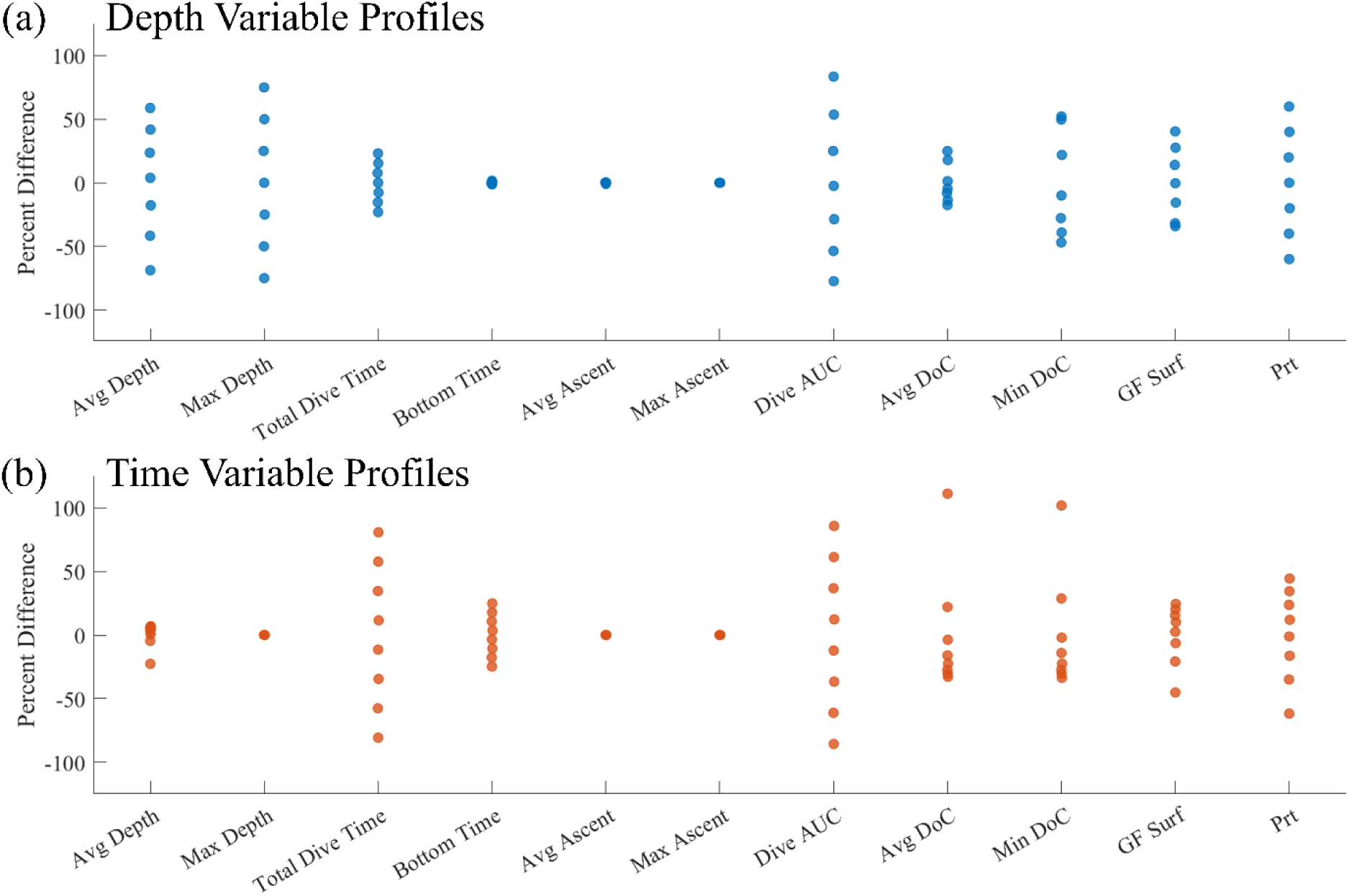
Percent difference of simulation dive profile metrics for (a) depth-variable and (b) time- variable simulations.

### Open water dives

The 40 field dives collected were found to have overlap in many cases (Fig. 6a). One diver reported difficulty with middle ear equilibration and was delayed in reaching the maximum depth, referred to as the ‘deviated profile’. Excluding the deviated profile from the field study data, an average depth of 17 meters and dive time 31 minutes was found (Fig. 6b,d). The maximum depth was 30.8 meters and the bottom time was 6.1 minutes (Fig. 6c,e). The average instantaneous ascent rate was 3.4 meters/minute, with average max of 15 meters/minute (Fig. 6f,g). The average AUC for the dives was 583 meters*minute (Fig. 6h). For the 39 dives, a GF_surf_ value of 0.81±0.008 while the deviated profile had a GF of 0.78 (Fig. 6i). Prt values of 10.0±0.31 and 14.1 was found for the 39 dives and deviated profile, respectively (Fig. 6j). An average RMSE of 0.0071±0.003 was calculated for the dives and 0.089 for the deviated profile (Fig. 6k).

**Figure 6.**
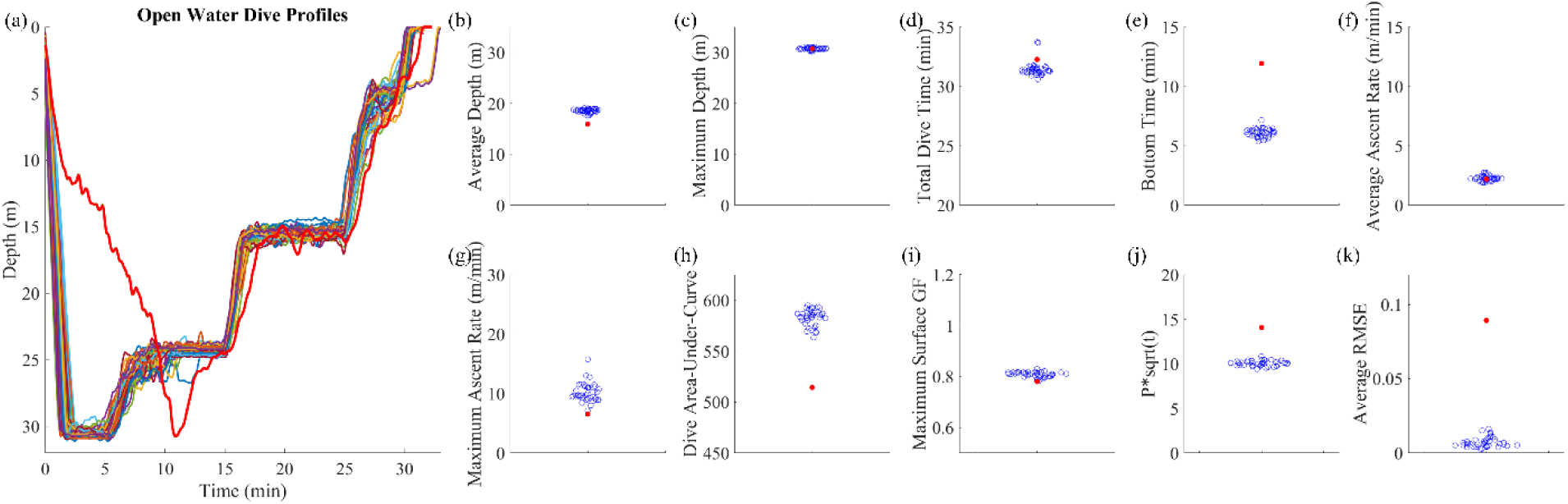
Field dive dataset with example metric calculations. The 40 dives are visualized in (a) with the following values: (b) Average Depth, (c) Maximum Depth, (d) Total Dive Time, (e) Bottom Time, (f) Average Ascent Rate, (g) Maximum Ascent Rate, (h) Dive Profile Area-Under-Curve, (i) Maximum Surface Gradient Factor (GF), (j) Maximum Pressure root Time (Prt), (k) Average Root-Mean-Square-Error (RMSE) of Bühlmann tissue compartments. In all panels, the red points represent the deviation profile, indicated by red line in (a).

A visual comparison of percent difference for each metric on the 40 field dives is presented in Figure 7. The deviated profile had the largest percent difference from the other 39 dives for bottom time, Prt, and average RMSE (Fig. 7a). Average RMSE percent difference for the deviated profile exceeded the reference value by 30-fold (Fig. 7b). Little to no difference between the 39 dives and deviated profile for average depth, maximum depth, total dive time, average ascent rate, maximum ascent rate, and AUC (Fig. 7a).

**Figure 7.**
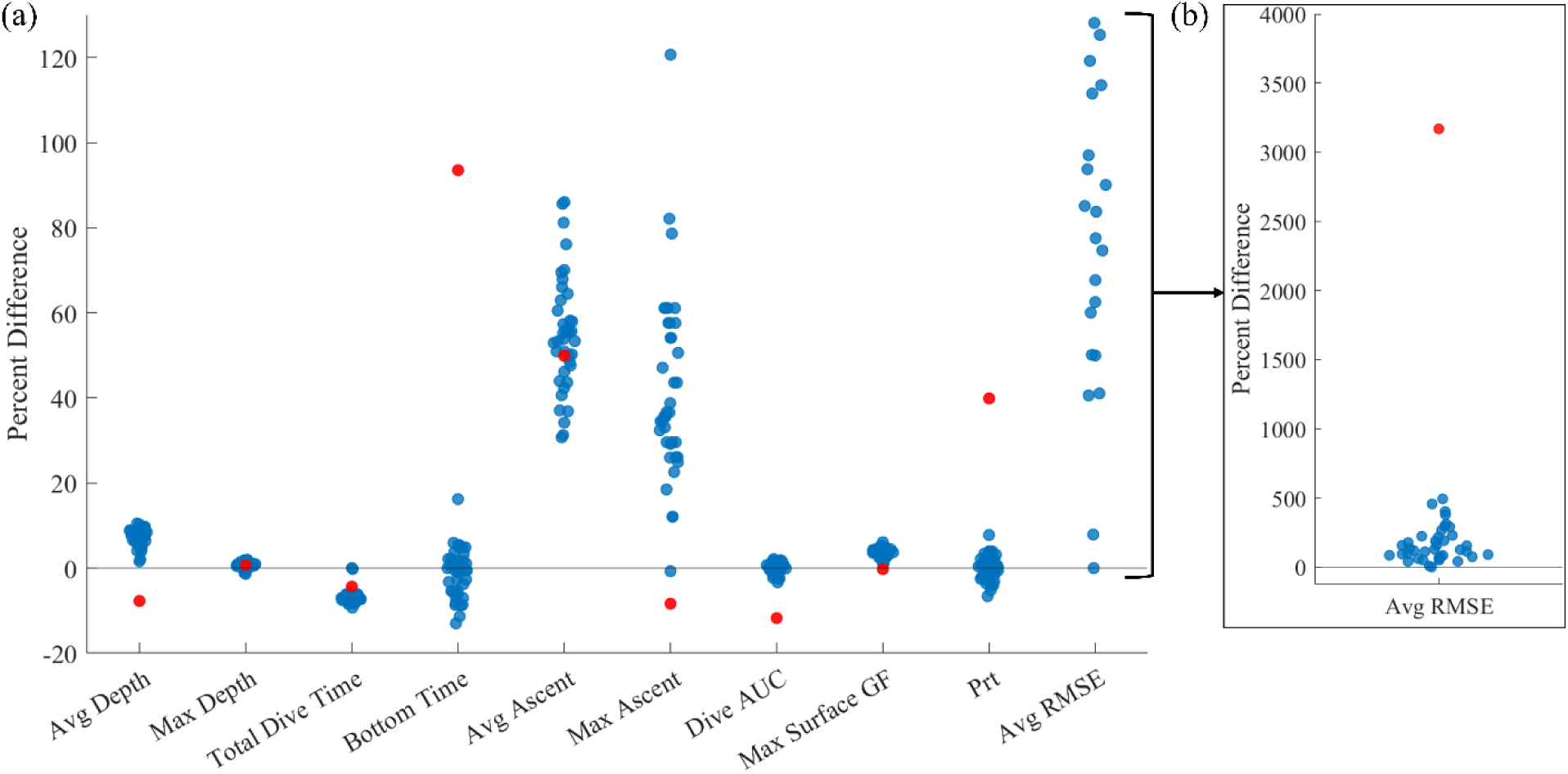
Percent difference of field dive profile metrics compared against the average from 39 consistent dive profiles. (a) Various dive metrics with (b) zoom out on Average Root-Mean-Square-Error (RMSE) of Bühlmann tissue compartments (note: for Average RMSE the minimum value from the 39 dives was used since the RMSE of reference profile = 0). For both panels, blue points represent 39 dive profiles and red indicates the deviated profile with slow time to reach maximum depth.

The subset of dives containing average of 39 field dives, deviation profile, and depth- augmented profile can be visualized in Fig. 8a. The minimum DoC value (compartment agnostic) is plotted over time in Fig. 8b. Additionally, a representation of the DoC across all 16-comparments is plotted at 5-minute intervals until 30-minutes after the end of the dive (Fig. 8c). Differences in minimum and average DoC are visible throughout the dive, although minimum DoC overlap occurs shortly after all dives were completed (Fig. 8b).

**Figure 8.**
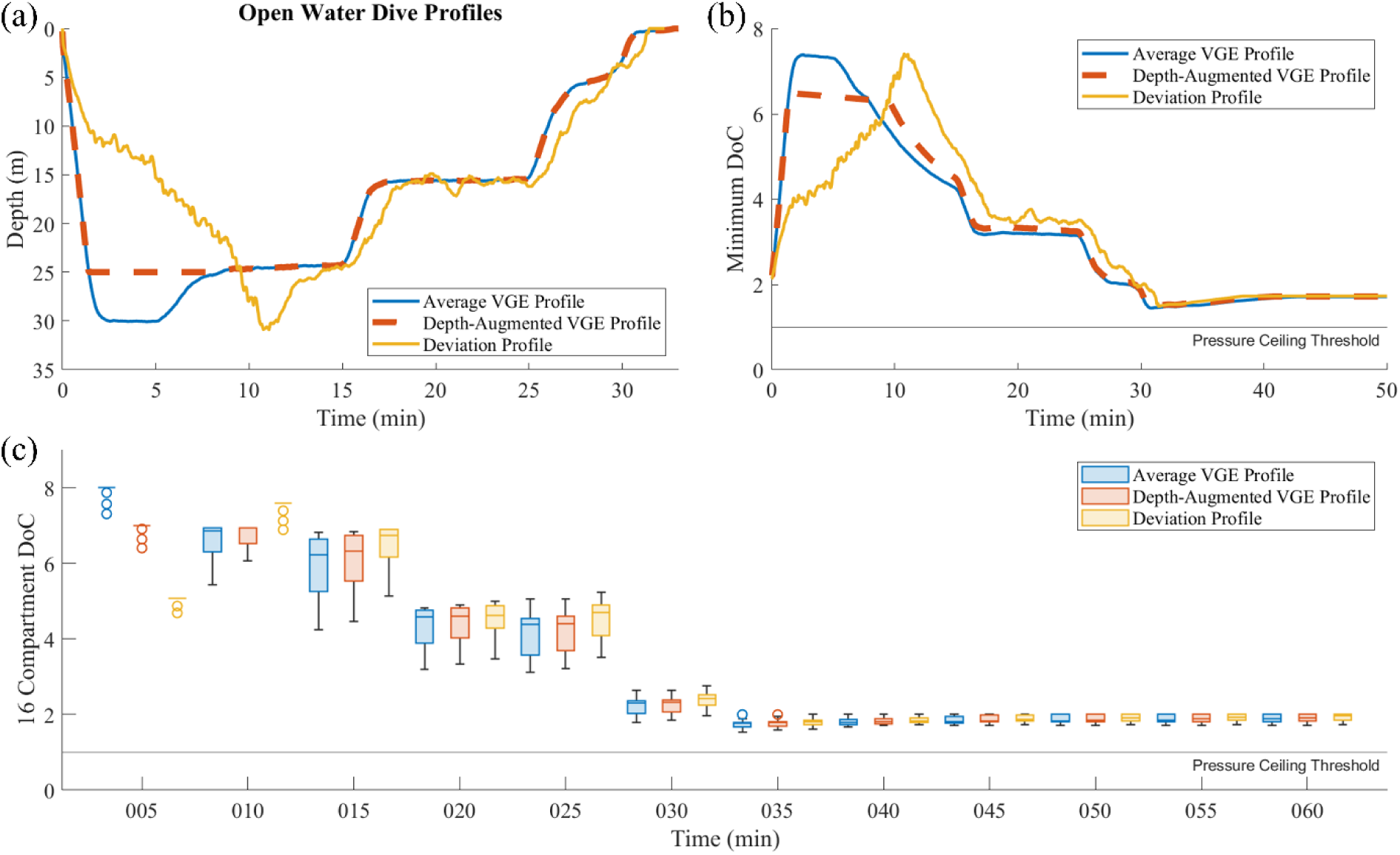
Example profiles with Degree of Conservatism (DoC) visulation. (a) Three dive profiles with a similar maximum depth or runtime. (b) Corresponding minimum DoC for each profile over time, extended after surfacing. (c) Evolution of DoC across the 16 Bühlmann tissue compartments throughout dive and after surfacing.

## Discussion

Decompression studies are not always conducted in hyperbaric chambers that simulate the underwater environment; instead, investigators often evaluate dives performed in open water (20, 21, 26, 31). While variability in VGE presence and DCS outcome has been described in both chamber and open-water dives (8, 20, 22), it is possible that minor individual differences in dive profile adherence may be a confounding variable for field studies. Previous work has described metrics for comparing dive profiles with large depth and time differences for human-dives (33, 36, 37) and animal studies (35, 43). Limited work has been conducted to evaluate minor deviations when study participants are attempting to follow a prescribed dive profile.

### Simulated Dive Profiles

Evaluation of the various metrics with the depth- and time-variable simulation profiles reiterates the contextual importance of the profiles being compared and the desired question. Unsurprisingly, the average and maximum depth calculations were sensitive to the depth-variable dataset, while total dive time and bottom time better differentiated the time-variable profiles. The average and maximum ascent rates were not sensitive to the depth nor time variations; however, for demonstration purposes, we set this value to be constant for all simulation profiles.

Notably, calculations of minimum DoC, maximum GF_surf_, and Prt ranked the dives in order of increasing depth or time for both datasets, confirming potential utility for dive risk estimation. These metrics tended to be more sensitive to both depth- and time-variable simulation datasets, compared to just depth or time only calculations. Prt ranked the longer duration profiles at a higher risk than the deeper dives, but this is likely an artifact of the depth ranges that were considered. However, the Prt calculation does not consider the decompression phase and would not be applicable for comparing multi-level dive profiles with a staged return to surface. Similar GF_surf_ values were found across the depth and time variations considered, demonstrating that increasing either will also increase the presumed risk of the dive. Due to similar theoretical inert gas totals, deeper dives with shorter bottom times produced a similar GF_surf_ to shallower dives with longer bottom times, a trait that underpins decompression modeling but complicates profile comparisons. Dive exposures are complex with a need for various depths and times depending on the objective, requiring a multi-faceted analysis approach.

### Open Water Dives

In this study, we proposed several methods to evaluate differences in theoretical inert gas tension under the Bühlmann framework and evaluated other common dive metrics on simulated profiles and open water human-dives measured using a dive computer.

Of the depth and time summary metrics, those with strongest sensitivity for detecting the deviation profile were average depth, bottom time, and Prt. The selected deviation profile had an extended time to reach the maximum depth and was delayed multiple minutes for the rest of the dive. This type of deviation could easily be captured by bottom time and Prt; however, because this metric excludes the decompression phase, it remains fundamentally inadequate for assessing overall dive profile similarity and associated decompression stress. The difference in maximum ascent rate was notable but may not be reliable given the time sampling rate (0.5 Hz) and could be attributed to arm movement or buoyancy adjustment. Average depth captures the net result over a period of time, without describing the overall dive path. The summary metrics offered quick evaluation of a given dive profile but appear insensitive to capturing the minor deviations of different dive profile attempts. Additionally, the simple depth and time-based calculations do not incorporate varied breathing gas mixes, preventing generalizability.

Expanding beyond summary measurements and leveraging DoC, we found the average RMSE comparing the individual attempts against the standard profile was most sensitive to the deviated profile. We derived percent difference to compare across the different units and scales; however, this may be biased for RMSE as the optimal value will approach zero and inflate the percent difference. Average RMSE demonstrated separation of the deviation profile from the rest in the magnitude calculations along. However, this approach is risk-agnostic and does not indicate if the dive was deeper or shallower, only describing the magnitude of deviation. While gradient factors were not used in the conventional sense for planning the decompression schedule, GF_surf_ was calculated for each compartment at the end of the dive. This metric represents how close the diver was to critical supersaturation and has been previously used to compare the risk level of a given dive profile (36). The GF_surf_ found similarity across all the field dives profiles and suggested that the deviated profile was safer than the average, differing from the Prt albeit with a smaller percent difference. This finding highlights the difficulty of evaluating different dive exposures for risk, predominantly that a standard calculation does not exist and existing methods produce conflicting results.

### Comparing Dive Profiles with Large Differences

Towards a generalizable metric to describe the risk level of a given dive profile, we ran a pilot test of several metrics against post-dive VGE outcome in real-world studies. The selected data was either from internally collected trials or previously published work (16, 44), all of which included post-dive VGE measurements following field standards (45). Ultrasound measurements of peak VGE presence are correlated with DCS risk, and an absence of VGE is highly indicative of diver safety (6, 18). Fig. 9 presents a visual of the dive profiles and comparison of the test metrics with the VGE outcome. The selected dive profiles varied in depth and bottom time, but all were conducted using air as the breathing gas. A potential trend was identified between the DoC based metric, calculated as number of minutes below a DoC threshold, where a longer time at risk corresponded with a lower peak VGE grade for the study. Aerobic exercise during a dive can augment post-dive VGE (11, 46–48), which is not currently taken into account by any metrics we evaluated. Comparison of dive profiles varying in depth, time, decompression technique, and environmental factors is not trivial and will require a multifaceted approach that extends beyond depth and time, potentially incorporating environmental and individual factors. Separately, Weinke previously proposed an end of dive risk estimator based on theoretical tissue supersaturation and total dive time, which has shown promise but not been externally validated (34). Future investigation into exercise during a dive is necessary to incorporate diver activity into dive profile risk evaluation.

**Figure 9.**
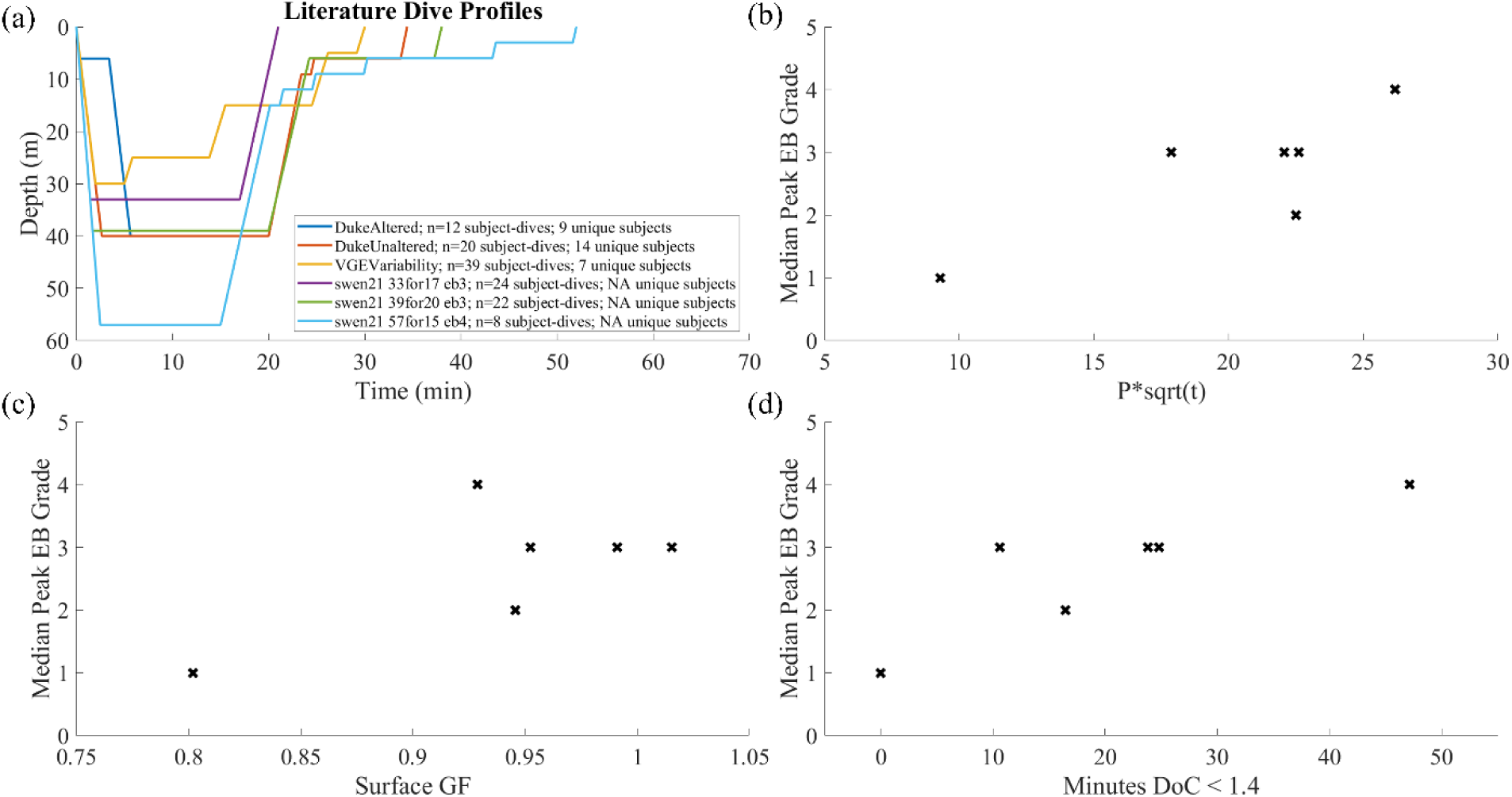
Preliminary comparison of dive profiles with larger deviations using a subset of metrics. (a) Six dive profiles from published work or internal datasets, all having reported peak VGE outcome. VGE outcome is presented as median peak EB grade within the reported group and is plotted against (b) Maximum Pressure root Time (Prt), (c) Maximum Surface Gradient Factor (GF), and (d) Total Time degree of conservatism (DoC) was below 1.4.

### Limitations

This work focused on previously published and validated decompression stress metrics (33, 37, 38, 41), as well as analysis under the Bühlmann ZHL-16C model (40). The ZHL-16C algorithm was selected for being nearly ubiquitous across dive computers. While design of the ZHL model compartments were informed by real tissue groups in the body with different perfusion rates, the 16 compartments are theoretical and cannot be extrapolated to represent anatomical locations. Furthermore, the ZHL-16C decompression framework assumes that respecting the maximum pressure limits will prevent bubble formation, hence maintaining the dissolved gas phase. However, circulating VGE are routinely identified in decompression trials, including dives within recreational limits, and DCS is known to occur despite obeying model thresholds (3). Previous work has even identified VGE presence during decompression with Doppler ultrasound (49), which is not measured or accounted for in current decompression models. Despite these limitations, the Bühlmann algorithm offers flexibility by incorporating depth, time, and breathing gas mix. The dives of interest in this study used air as a breathing gas, yet by adjusting the inert gas half-times by a multiple of 2.65 the model can accommodate helium kinetics (40), which was not explored in this work. Future analysis could include dives relying on breathing mixes with helium as a component or utilizing an oxygen-rich decompression gas mix.

## Conclusion

DCS remains the primary limiting factor in military and commercial diving operations. Many studies have previously demonstrated variability in DCS outcome and VGE presence despite having study participants conduct the same prescribed dive in open water. However, the impact of minor profile deviations throughout the dive has not been evaluated as a contributing factor to variability. We have developed a framework under the Bühlmann algorithm to measure dive profile consistency (or lack thereof) across attempts at a prescribed dive. The framework and other common dive risk evaluation metrics were tested using both simulated and open-water dive profiles, demonstrating differences in sensitivity to dive profile deviations. Quantitative analysis of dive profiles and *a priori* establishment of deviation tolerance could produce more reliable experimental data, and the dive risk metrics may be compared against VGE or DCS outcome in future studies.

## Acknowledgements

We gratefully acknowledge the Divers Alert Network, and particularly Rhiannon Brenner, for assistance with collection of the open-water dive profiles used as illustrative examples in this manuscript.

This work was supported by the Divers Alert Network (DAN) research foundation through grants #DAN-UNC-1 and DAN-UNC-2 to Virginie Papadopoulou (V.P.). Frauke Tillmans (F.T.), Vice President of Research at DAN, is a co-author on this manuscript. All scientific decisions, including study design, data analysis, interpretation, manuscript preparation, and the decision to publish, were made by the authors. DAN did not exercise authority over the conduct of the research or publication of the results beyond the scientific contributions of F.T. in her capacity as a co-author.

## Notes

### Competing Interest Statement

The authors have declared no competing interest.

## References

1. Mitchell SJ. Decompression illness: a comprehensive overview. Diving Hyperb Med 54: 1–53, 2024.

2. Mitchell SJ, Bennett MH, Moon RE. Decompression Sickness and Arterial Gas Embolism. New Engl J Med 386: 1254–1264, 2022.

3. Tillmans F. DAN Annual Diving Report 2020 Edition: A report on 2018 diving fatalities, injuries, and incidents. Durham, NC: 2021.

4. Doolette DJ, Gault KA, Gutvik CR. Sample size requirement for comparison of decompression outcomes using ultrasonically detected venous gas emboli (VGE): Power calculations using Monte Carlo resampling from real data. Diving Hyperb Med 44: 14–19, 2014.

5. Blogg SL, Lang MA, Møllerløkken A. Validation of Dive Computers. In: Diving for Science 2012, edited by Steller D, Lobel L. AAUS, 2012, p. 62–81.

6. Currens JB, Doolette DJ, Murphy FG. Venous gas emboli (VGE) in 2-D echocardiographic images following movement: grading and association with cumulative incidence of decompression sickness. Diving Hyperb Med 54: 44–50, 2025.

7. Doolette D. Venous gas emboli detected by two-dimensional echocardiography are an imperfect surrogate endpoint for decompression sickness. Diving Hyperb Med 46: 4–10, 2016.

8. Hess HW, Wheelock CE, James ES, Stooks JL, Clemency BM, Hostler D. Variability in venous gas emboli following the same dive at 3,658 meters. Undersea Hyperb Med 48: 469–476, 2021.

9. Nishi RY. Design of Decompression Trials - DCIEM Experience. In: Proceedings of Repetitive Diving Workshop, edited by Lang MA, Vann RD. Durham: American Academy of Underwater Sciences, 1991, p. 311–320.

10. Brett KD, Nugent NZ, Fraser NK, Bhopale VM, Yang M, Thom SR. Microparticle and interleukin-1β production with human simulated compressed air diving. Sci Rep 9, 2019.

11. Jankowski LW, Tikuisis P, Nishi RY. Exercise effects during diving and decompression on postdive venous gas emboli. Aviat Space Environ Med 75: 489–495, 2004.

12. Doolette D, Gerth W, Gault K. Redistribution of decompression stop time from shallow to deep stops increases incidence of decompression sickness in air decompression dives, NEDU TR 11-06. Panama City, FL, USA: 2011.

13. Doolette DJ, Gault KA, Gerth WA. Manipulating Frequency or Duration of Air Breaks During Oxygen Decompression Did Not Reveal a Direct Contribution of Oxygen to Decompression Stress, NEDU TR 17-13. Panama City, FL, USA: 2017.

14. Gerth WA, Ruterbusch VL, Long ET. The influence of thermal exposure on diver susceptibility to decompression sickness, NEDU TR 03-09. Panama City, FL, USA: 2007.

15. Andrew BT, Doolette DJ. Manned validation of a US Navy Diving Manual, Revision 7, VVal-79 schedule for short bottom time, deep air decompression diving. Diving Hyperb Med 50: 43, 2020.

16. Hjelte C, Plogmark O, Silvanius M, Ekstrom M, Franberg O. Risk assessment of SWEN21 a suggested new dive table for the Swedish armed forces: bubble grades by ultrasonography. Diving Hyperb Med 53: 299–305, 2023.

17. Gennser M, Jurd KM, Blogg SL. Pre-dive exercise and post-dive evolution of venous gas emboli. Aviat Space Environ Med 83: 30–34, 2012.

18. Sawatzky KD. The relationship between intravascular Doppler-detected gas bubbles and decompression sickness after bounce diving in humans, MS Thesis. York University: 1991.

19. Møllerløkken A, Breskovic T, Palada I, Valic Z, Dujić Ž, Brubakk AO. Observation of increased venous gas emboli after wet dives compared to dry dives. Diving Hyperb Med 41: 124–128, 2011.

20. Papadopoulou V, Germonpré P, Cosgrove D, Eckersley RJ, Dayton PA, Obeid G, Boutros A, Tang MX, Theunissen S, Balestra C. Variability in circulating gas emboli after a same scuba diving exposure. Eur J Appl Physiol 118: 1255–1264, 2018.

21. Carturan D, Boussuges A, Vanuxem P, Bar-Hen A, Burnet H, Gardette B. Ascent rate, age, maximal oxygen uptake, adiposity, and circulating venous bubbles after diving. J Appl Physiol 93: 1349–1356, 2002.

22. Doolette DJ, Murphy FG. Within-diver variability in venous gas emboli (VGE) following repeated dives. Diving Hyperb Med 53: 333–339, 2023.

23. Thom SR, Milovanova TN, Bogush M, Yang M, Bhopale VM, Pollock NW, Ljubkovic M, Denoble P, Madden D, Lozo M. Bubbles, microparticles, and neutrophil activation: changes with exercise level and breathing gas during open-water SCUBA diving. J Appl Physiol 114: 1396–1405, 2013.

24. Zanchi J, Ljubkovic M, Denoble PJ, Dujic Z, Ranapurwala S, Pollock NW. Influence of repeated daily diving on decompression stress. Int J Sports Med 35: 465–468, 2014.

25. Susilovic-Grabovac Z, Obad A, Duplančić D, Banić I, Brusoni D, Agostoni P, Vuković I, Dujic Z, Bakovic D. 2D speckle tracking echocardiography of the right ventricle free wall in SCUBA divers after single open sea dive. Clin Exp Pharmacol Physiol 45: 234–240, 2018.

26. Ljubkovic M, Marinovic J, Obad A, Breskovic T, Gaustad SE, Dujic Z. High incidence of venous and arterial gas emboli at rest after trimix diving without protocol violations. J Appl Physiol 109: 1670–1674, 2010.

27. Ljubkovic M, Dujic Z, MØllerlØkken A, Bakovic D, Obad A, Breskovic T, Brubakk AO. Venous and arterial bubbles at rest after no-decompression air dives. Med Sci Sports Exerc 43: 990–995, 2011.

28. Bradbury KE, DiMarco KG, Futral JE, Lord RN, Edward JA, Barak O, Glavičić I, Miloš I, Drvis I, Dujić Ž. The maintenance of core temperature in SCUBA divers: Contributions of anthropometrics, patent foramen ovale, and non-shivering thermogenesis. J Sci Med Sport 27: 820–827, 2024.

29. Lautridou J, Pichereau V, Artigaud S, Bernay B, Barak O, Hoiland R, Lovering AT, Eftedal I, Dujic Z, Guerrero F. Evolution of the plasma proteome of divers before and after a single SCUBA dive. Proteomics Clin Appl 11: 1700016, 2017.

30. Lambrechts K, Pontier JM, Balestra C, Mazur A, Wang Q, Buzzacott P, Theron M, Mansourati J, Guerrero F. Effect of a single, open-sea, air scuba dive on human micro- and macrovascular function. Eur J Appl Physiol 113: 2637–2645, 2013.

31. Dujić Ž, Palada I, Obad A, Duplančić D, Baković D, Valic Z. Exercise during a 3-min decompression stop reduces postdive venous gas bubbles. Med Sci Sports Exerc 37: 1319– 1323, 2005.

32. Wienke BR. Diving decompression models and bubble metrics: modern computer syntheses. Comput Biol Med 39: 309–331, 2009.

33. Baker EC. Understanding M-values. Immersed 3: 23–27, 1998.

34. Wienke BR. Dive Computer Profile Data and on the Fly and End of Dive Risk Estimators. Journal of Applied Biotechnology & Bioengineering 5, 2018.

35. Buzzacott P, Lambrechts K, Mazur A, Wang Q, Papadopoulou V, Theron M, Balestra C, Guerrero F. A ternary model of decompression sickness in rats. Comput Biol Med 55: 74–78, 2014.

36. Cialoni D, Pieri M, Balestra C, Marroni A. Dive risk factors, Gas Bubble formation, and decompression illness in recreational SCUBA diving: Analysis of DAN Europe DSL data base. Front Psychol 8, 2017.

37. Hempleman H, Crocker W, Taylor H. Investigation into the decompression tables, Report III, Part A: A New Theoretical Basis for the Calculation of Decompression Tables. 1952.

38. Shields TG, Duff PM, Wilcock SE, Giles R. Decompression sickness from commercial offshore air-diving operations on the UK continental shelf during 1982 to 1988. Subtech 23: 259–77, 1989.

39. Weathersby P, Homer L, Flynn E. On the likelihood of decompression sickness. J Appl Physiol 57: 815–825, 1984.

40. Bühlmann AA. Decompression—decompression sickness. Berlin: Springer Science & Business Media, 1984.

41. Papadopoulou V. Computational and experimental techniques towards optimising the cardiovascular risk assessment of hyperbaric decompression stress caused by circulatory bubble dynamics, PhD Thesis. Imperial College London: 2015.

42. United States ND. US Navy Diving Manual, Rev. 7, Volume 9: The Air Decompression Table. 2017.

43. Lillo RS, Parker EC. Mixed-gas model for predicting dcs in rats. J Appl Physiol 89: 2107– 2116, 2000.

44. Currens JB, Natoli MJ, Eltz KM, Morales G, Bautista KJB, Dayton PA, Lance RM, Oralkan O, Yamaner FY, Moon RE, Papadopoulou V. Human decompression in real time: programmable ultrasound imaging during hyperbaric exposure. bioRxiv [preprint] 2026.07.22.737513, 2026.

45. Møllerløkken A, Blogg SL, Doolette DJ, Nishi RY, Pollock NW. Consensus Development Conference: Consensus guidelines for the use of ultrasound for diving research. Diving Hyperb Med 46, 2016.

46. Jauchem JR. Effects of exercise on the incidence of decompression sickness: a review of pertinent literature and current concepts. Int Arch Occup Environ Health 60: 313–319, 1988.

47. Dujić Ž, Duplančic D, Marinovic-Terzić I, Baković D, Ivančev V, Valic Z, Eterović D, Petri NM, Wisløff U, Brubakk AO. Aerobic exercise before diving reduces venous gas bubble formation in humans. Journal of Physiology 555: 637–642, 2004.

48. Vann RD, Butler FK, Mitchell SJ, Moon RE. Decompression illness. The Lancet 377: 153–164, 2011.

49. Neuman TS, Hall DA, Linaweaver Jr PG. Gas phase separation during decompression in man: ultrasound monitoring. Undersea Biomed Res 3: 121–130, 1976.

